# The EGF-motif containing protein SPE-36 is a secreted protein required for sperm function at fertilization in *C. elegans*

**DOI:** 10.1101/2021.07.07.451551

**Authors:** Amber R. Krauchunas, Matthew R. Marcello, A’Maya Looper, Xue Mei, Emily Putiri, Gunasekaran Singaravelu, Iqra I. Ahmed, Andrew Singson

## Abstract

The growing number of genes specifically required for fertilization suggests that there is a significant amount of molecular complexity at the sperm-egg interface. Thus, we have adopted a model of a “fertilization synapse” where specialized zones of interaction and multi-protein complexes mediate gamete interaction and fusion. The fertilization synapse is likely to be composed of both *trans* and *cis* protein-protein interactions at the surface of each gamete. Mutations in the *Caenorhabditis elegans spe-36* gene result in a sperm-specific fertility defect. Surprisingly, *spe-36* encodes a secreted EGF-motif containing protein that functions cell autonomously. Despite the fact that morphology and migratory behavior of *spe-36* sperm are indistinguishable from wild-type sperm, *spe-36* sperm make close contact with oocytes but fail to fertilize them. The genetic requirement for a secreted sperm-derived protein for fertilization is novel and represents a paradigm-shifting discovery in the molecular understanding of fertilization.

## Introduction

Fertilization is a complex process and recent work indicates the regulation of this molecular event is even more complicated than previously thought. In mammals, we know of a single protein-protein interaction between the gametes: binding of the egg surface protein JUNO to the sperm surface protein IZUMO1 (Aydin et al., 2016; Bianchi et al., 2014; Inoue et al., 2005; Jean et al., 2019; Ohto et al., 2016). While the JUNO-IZUMO1 interaction is necessary for fertilization, it is not sufficient for cellular fusion (Bianchi et al., 2014; Chalbi et al., 2014). As such, we clearly do not yet know all of the protein interactions necessary for adhesion and fusion between the two gametes. Until we have identified all of the genes and proteins involved, our ability to understand the *trans*, and *cis*, protein interactions that mediate fertilization will be limited.

Forward genetic screens provide the opportunity to discover novel genes necessary for fertilization. The nematode *C. elegans* is an excellent model system for such studies (Singson, 2001). Identified through forward genetics, the *spe-9* gene was the first gene to be identified in *C. elegans* as necessary for fertilization (Singson et al., 1998). Since then, genetic screens in *C. elegans* have led to the identification of nine additional sperm genes necessary for fertilization [including Mei *et al*., co-submission and the *spe-36* gene reported in this paper](Chatterjee et al., 2005; Kroft et al., 2005; L’Hernault et al., 1988; Roberts and Ward, 1982; Singaravelu et al., 2015; Wilson et al., 2018; Xu and Sternberg, 2003). This includes *spe-45*, which encodes an immunoglobulin containing protein similar to the mammalian protein IZUMO1 (Krauchunas and Singson, 2016; Nishimura et al., 2015; Singaravelu et al., 2015). Mutations in any one of these genes results in healthy adult animals that are sterile. Sperm from these mutants have normal morphology, migrate to and maintain their position at the site of fertilization in the reproductive tract, and make contact with eggs but fail to fertilize the eggs. This same phenotype is observed in mammals lacking *Izumo1, Spaca6, Tmem95, Sof1*, or *FIMP* (Barbaux et al., 2020; Fujihara et al., 2020; Inoue et al., 2005; Lamas-Toranzo et al., 2020; Lorenzetti et al., 2014; Noda et al., 2020). The growing number of genes required for fertilization has shifted our view from a ligand-receptor pair to the model of a fertilization synapse. In this model specialized zones of interaction and multi-protein complexes composed of both *trans* and *cis* protein-protein interactions mediate gamete interaction and fusion.

Here we report the novel discovery of SPE-36 as a sperm-derived secreted protein that is necessary for fertilization. Sperm from *spe-36* mutants look phenotypically normal, are motile, and can migrate to the site of fertilization. However, sperm that do not produce SPE-36 protein cannot fertilize. SPE-36 and SPE-51 (co-submission Mei *et al*.) are the first secreted proteins found to be necessary for fertilization. Discovery of these secreted proteins sheds new light on the complex nature of fertilization and supports the model of a fertilization synapse.

## Materials and methods

### Strains and culture methods

General maintenance and crosses of *C. elegans* were performed as described previously (Brenner, 1974). Worms were cultured on MYOB (Modified Youngren’s, Only Bacto-peptone) plates seeded with OP50 *E. coli*, with the exception of *spe-36(as1)* fertility assays which were performed on NGM plates. The *spe-36(it114*) allele comes from an ethyl methanesulfonate (EMS) mutagenesis screen for genes with fertility defects performed by Dr. Diane Shakes while in the Kemphues laboratory at Cornell University. The *spe-36(as1)* and *spe-36(as6)* alleles were identified through a F1 non-complementation screen with *spe-36(it114)* performed in the Singson laboratory. Other strains used: Bristol N2, CB4856, *him-5(e1490), fem-1(hc17)*, CB61 *dpy-5(e61)*, MDX44 *cylc-2(mon2[cylc-2::mNG^3xFLAG])*, SL438 *spe-9(eb19)* I; *him-5(e1490)* V; *ebEx126*, RT36 *arIs37 [myo-3p::ssGFP]*, HCL67 *unc-119(ed3)* III; *uocIs1 [eft-3p::Cas9(dpiRNA)::tbb-2 3’ UTR + unc-119(+)]*. Strains created in this study: ARK5 *spe-36(as6); asEx96 [PCR product of genomic spe-36]*, ARK6 *spe-36(as6); him-5(e1490); asEx96, cylc-2(mon2); spe-36(as6); him-5(e1490), myo-3p::spe-36::gfp, myo-3p::spe-9::gfp*, ARK7 *spe-36(nwk1[spe-36::gfp]); him-5(e1490)*.

### Fertility Assays

The number of self-progeny for wild type (N2), *spe-36* mutants, and the transgenic rescue stocks was determined by placing single hermaphrodites on separate culture plates and culturing at 20°C. Worms were transferred to fresh plates every 24 hours until each worm stopped producing progeny. After the worm was transferred off the plate the number of unfertilized oocytes on the plate was counted. The number of progeny on each plate was counted 3 days after the parent was removed. To determine male fertility, and fertility of *spe-36* hermaphrodites mated to fertile males, crosses were set up in a ratio of 1 hermphrodite:4 males. Males were removed after 24 hours and hermaphrodites were transferred to fresh plates every day for 3 days. The number of progeny on each plate was counted 3 days after the parent was removed.

### Fertility assays to determine cell autonomy of SPE-36

*fem-1(hc17)* animals were reared at 25°C until the L4 stage. All other strains were cultured at 20°C and all crosses were performed at 20°C. *spe-36(as6); him-5* males were crossed with either *fem-1(hc17)* or *dpy-5(e61)* hermaphrodites in a ratio of 4:1. Alternatively, *spe-36(as6)* hermaphrodites were crossed with either *spe-9(eb19); him-5* or *spe-36(as6); him-5* males in a ratio of 1:3. For all crosses, parents were removed from the plate after 24 hours and the number of progeny produced in that 24 hour period was counted 3 days later.

### Phenotypic analysis

Phenotypic analysis of *spe-36* mutants was largely conducted as described previously (Marcello et al., 2013). *spe-36* animals lacking the rescuing transgene were identified as failing to express the *myo-3p::GFP* reporter in either their somatic tissue or progeny. Light microscopy, DAPI staining, sperm isolation and *in vitro* activation with Pronase were all performed as described in Krauchunas et al.(2018).

### Electron microscopy

Transmission Electron Microscopy (TEM) of sperm in the spermatheca were obtained as previously described. Worms were anesthetized with 0.5% propylene phenoxytol in M9 buffer and then fixed in a glass dish with 2.5% gluteraldehyde in 0.1 M HEPES buffer for 3–4 hours. The worms were then post-fixed in 1% osmium tetroxide for 1 hour, after which they were aligned on a thin layer of 1% agar. They were then covered with a drop of molten 1% agar and worm-containing blocks of agar cut out. These blocks were dehydrated in a graded series of acetone and infiltrated and imbedded in Epon–Spurr resin. Thin sections (90–100nm) were cut using a Reichert Ultracut E microtome with a diamond knife. The sections were lifted to copper grids and double-stained using ethanoic uranyl acetate (10 min) and lead citrate (2 min). Sections were examined and photographed at 80 kV with a JEOL 1200 electron microscope.

### Assessing mating, sperm transfer, and migration

*cylc-2(mon2[C41G7*.*6::mNG^3xFLAG)* was crossed into *spe-36(as6); him-5(e1490)* to produce *cylc-2(mon2[C41G7*.*6::mNG^3xFLAG) I; spe-36(as6) IV; him-5(e1490) V*. Wild-type and *spe-36* males with mNG marked sperm were crossed with L4 N2 hermaphrodites in a ratio of 10 hermaphrodites to 30 males per plate (3 plates per group). Approximately 24 hours later the males were removed from the plates. 4-6 hours after the males were removed the hermaphrodites were mounted on 2% agarose pads in M9 with levamisole and imaged.

### Whole genome sequencing and analysis

*spe-36(it114)* hermaphrodites and *spe-36(as6)* hermaphrodites were crossed with males from the polymorphic wild-type Hawaiian strain CB4856. For each allele, individual F2 hermaphrodites were picked at the L4 stage to 20 24-well plates and scored the following day for sterility. Approximately 100 sterile (*spe-36* homozygous) hermaphrodites for each allele were pooled separately for preparation of genomic DNA as described in Smith et al. (2016). 10 ng of fragmented DNA was used for library preparation using KAPA Hyper Prep Kit (KAPA Biosystems) according to the manufacturer’s instruction. PCR was performed on library DNA for 12 cycles after which all libraries were pooled according to Illumina HiSeq specifications. Sequencing was performed at JHU Genetic Resources Core Facility using an Illumina HiSeq 2500 platform with a 50-nt single-end run and dedicated index sequencing. Demultiplexed data was analyzed using MiModD (https://mimodd.readthedocs.io/en/latest/nacreousmap.html).

### Preparation of transgenic rescue lines

Young adult N2 hermaphrodites were microinjected with a PCR product of genomic F40F11.4 including 479 bp of DNA upstream and 303 bp downstream of the coding sequence along with myo-3p::gfp (100 μg/ml) as a transformation marker. Once transgenic lines were established the extrachromosomal array was crossed into *spe-36(as6)* to establish rescue of the sterile phenotype.

### Preparation of transgenic lines expressing *spe-36*::*gfp* or *spe-9::gfp* in muscle

*spe-36* and *spe-9* cDNAs were PCR amplified with primers designed for Gibson assembly and containing sequence overlap with the plasmid pJF25 (myo-3p::ssGFP, gift from Barth Grant). Restriction digest was performed to linearize pJF25 and remove the signal peptide upstream of GFP. Gibson assembly was used to insert either *spe-36* cDNA or *spe-9* cDNA in frame between the myo-3 promoter and GFP protein sequence. Plasmids were microinjected into N2 worms along with myo-2p::mCherry as a transformation marker. Once transgenic lines were established worms were imaged using a Zeiss Universal microscope to confirm expression of GFP in the body wall muscles and determine if the GFP signal was also present in the coelomocytes.

### CRISPR editing

CRISPR editing was carried out similar to the protocol described in (Paix et al., 2017). An sgRNA targeting *spe-36* was ordered from Synthego with the sequence AUUCGCAUUGUCUGAAGAGA. A repair template containing homology arms to the endogenous *spe-36* locus flanking a gfp coding sequence that was engineered to be resistant to piRNA silencing (Zhang et al., 2018) was synthesized as a gene fragment by IDT. CRISPR-Cas9 ribonucleoprotein complexes and repair templates were injected into HCL67 hermaphrodites which express Cas9 in the germline (Zhang et al., 2018). Edited worms were sequenced by Sanger sequencing to confirm the gfp insertion. Worms were then outcrossed to *him-5(e1490)* and genotyped by PCR to confirm they were homozygous for the gfp insertion and no longer carrying the Cas9 transgene.

### Immunostaining

Hermaphrodites were dissected and immunostained similar to previously described methods (Arduengo et al., 1998; Zannoni et al., 2003). Approximately 20 L4 stage hermaphrodites and 25 L4 males were transferred onto the same plate and left for 24 hours. The following day, hermaphrodites were removed and dissected using a needle to release sperm onto charged adhesive slides. Samples were fixed with 4% paraformaldehyde in 1X sperm media + dextrose. Following fixation and permeabilization, slides were blocked with 20% Normal Goat Serum and 0.9% Bovine Serum Albumin in PBS. Polyclonal rabbit anti-GFP primary antibody was used at a 1:500 dilution in PBS (Novus Biologicals, Littleton, CO). Alexa Fluor 488-conjugated goat anti-Rabbit IgG secondary antibody was used at a 1:200 dilution in PBS (Thermo FisherScientific, Waltham, MA). After the final wash, antifade mounting medium with DAPI (Vector- Laboratories, Burlingame, CA) was added to the slides and samples were covered with a coverslip. Slides were imaged using a ZEISS Axio Observer D1 inverted microscope using a 100X objective.

## Results

### *spe-36* mutants are sterile due to a sperm-specific defect

Unmated *spe-36* hermaphrodites produce little to no progeny despite having a high number of ovulations (Figure 1A and S1). However, *spe-36* hermaphrodites produce viable progeny when crossed to wild-type (N2) males, indicating that *spe-36* oocytes and the somatic gonad are functional. Because *spe-36* hermaphrodites display no additional mutant phenotypes, we conclude that these hermaphrodites have a sperm-specific defect. To assess male fertility, we compared the ability of *spe-36* and wild-type males to sire outcross progeny from feminized (*fem-1*) hermaphrodites that do not produce any self sperm. In sharp contrast to wild-type controls, *spe-36* males sired a greatly reduced number of outcross progeny (Figure 1A). Therefore, the *spe-36* mutant defect affects both male and hermaphrodite sperm.

**Figure 1.**
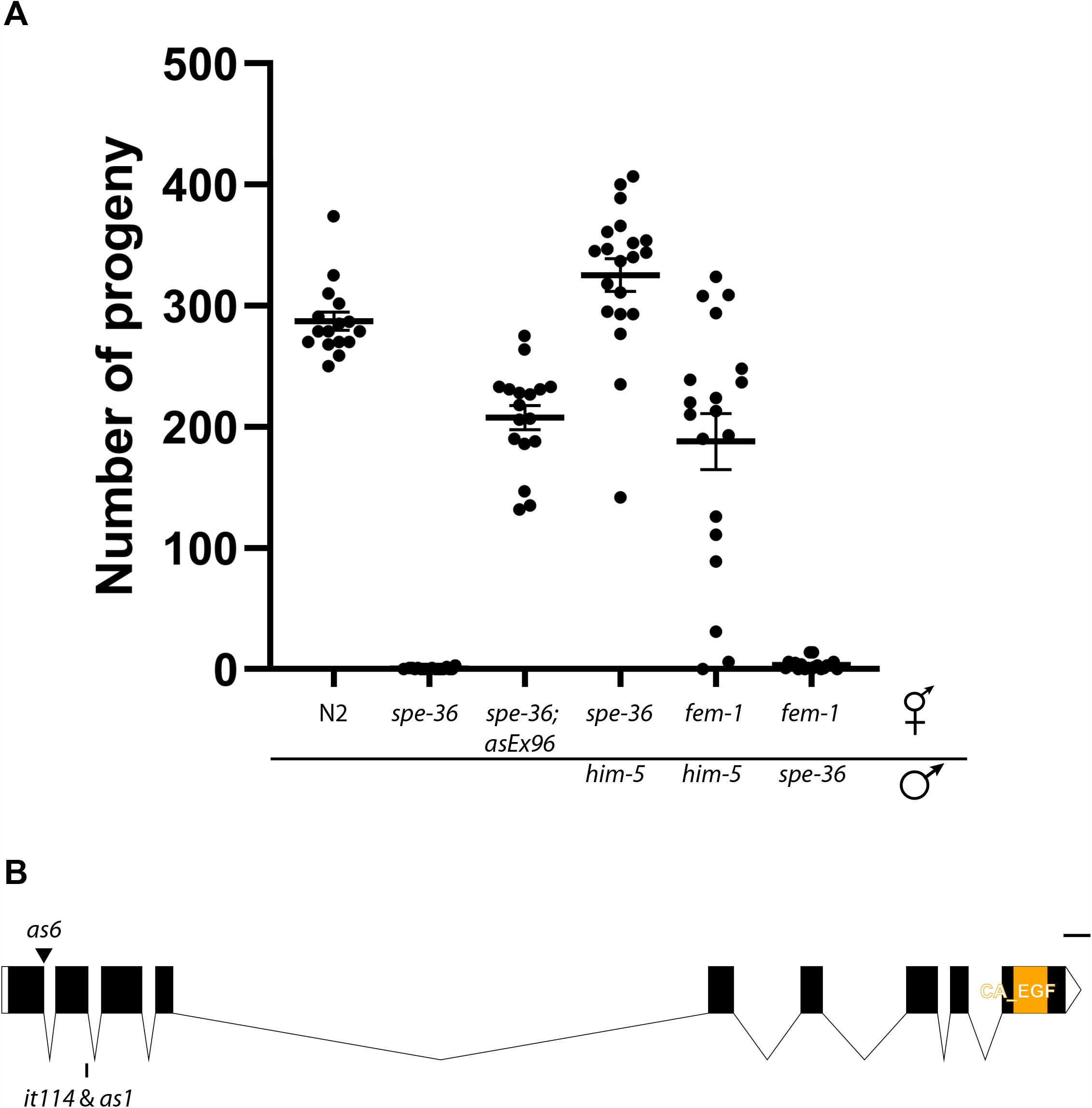
*spe-36* mutants are sterile due to a sperm defect. (A) Hermaphrodite and male fertility was determined for *spe-36(as6)* mutants. Unmated *spe-36(as6)* hermaphrodites produce no self-progeny (avg = 0.6 progeny, n = 19) and an extrachromosomal array containing a PCR product of *F40F11*.*4* (*asEx96*) restores fertility to *spe-36(as6)* hermaphrodites. *spe-36(as6)* hermaphrodites also produce progeny when mated to wild-type males, showing that the sterility is caused by a defect in *spe-36(as6)* sperm. When *spe-36(as6)* males are mated to feminized hermaphrodites they fail to produce progeny, indicating that *spe-36* is also necessary for male fertility. Error bars indicate standard error. (B) Schematic of the *spe-36* gene structure including the locations of the *it114, as1*, and *as6* mutations and the region that encodes the EGF motif (amino acids 282 – 329). The *it114* and *as1* alleles have the same nucleotide change at position chrIV: 11,591,836 resulting in a C -> Y change at amino acid 104. *as6* has a nucleotide change at position chrIV: 11,592,109 that alters the splice donor site of the first intron. (C. elegans Feb. 2013 (WBcel235/ce11) Assembly) Scale bar represents 100 bp.

Direct observation of *spe-36* hermaphrodite reproductive tracts confirmed the presence of oocytes and sperm. However, instead of developing embryos in the uterus we observed unshelled eggs that appear similar to the unfertilized oocytes in the oviduct (Figure 2A). DAPI staining to observe the DNA content in these cells showed a single DNA mass in each egg (Figure 2B). Since the centrioles are supplied to the egg by the sperm, a single DNA mass in each egg in the uterus indicates that no sperm entered these eggs (Albertson, 1984; Ward and Carrel, 1979). Yet we know that the sperm must have contacted the eggs when the eggs moved through the spermatheca to enter the uterus, as sperm are clearly present in the spermatheca. Finally, we find that sperm are still present in the spermathecae of older *spe-36* adult hermaphrodites (Figure 2D). This is in contrast to wild-type hermaphrodites where successful fertilization depletes all self sperm in the spermathecae by day 4 of adulthood (Figure 2C). From these data we conclude that *spe-36* mutant sperm are incapable of fertilizing either *spe-36* or wild-type oocytes.

**Figure 2.**
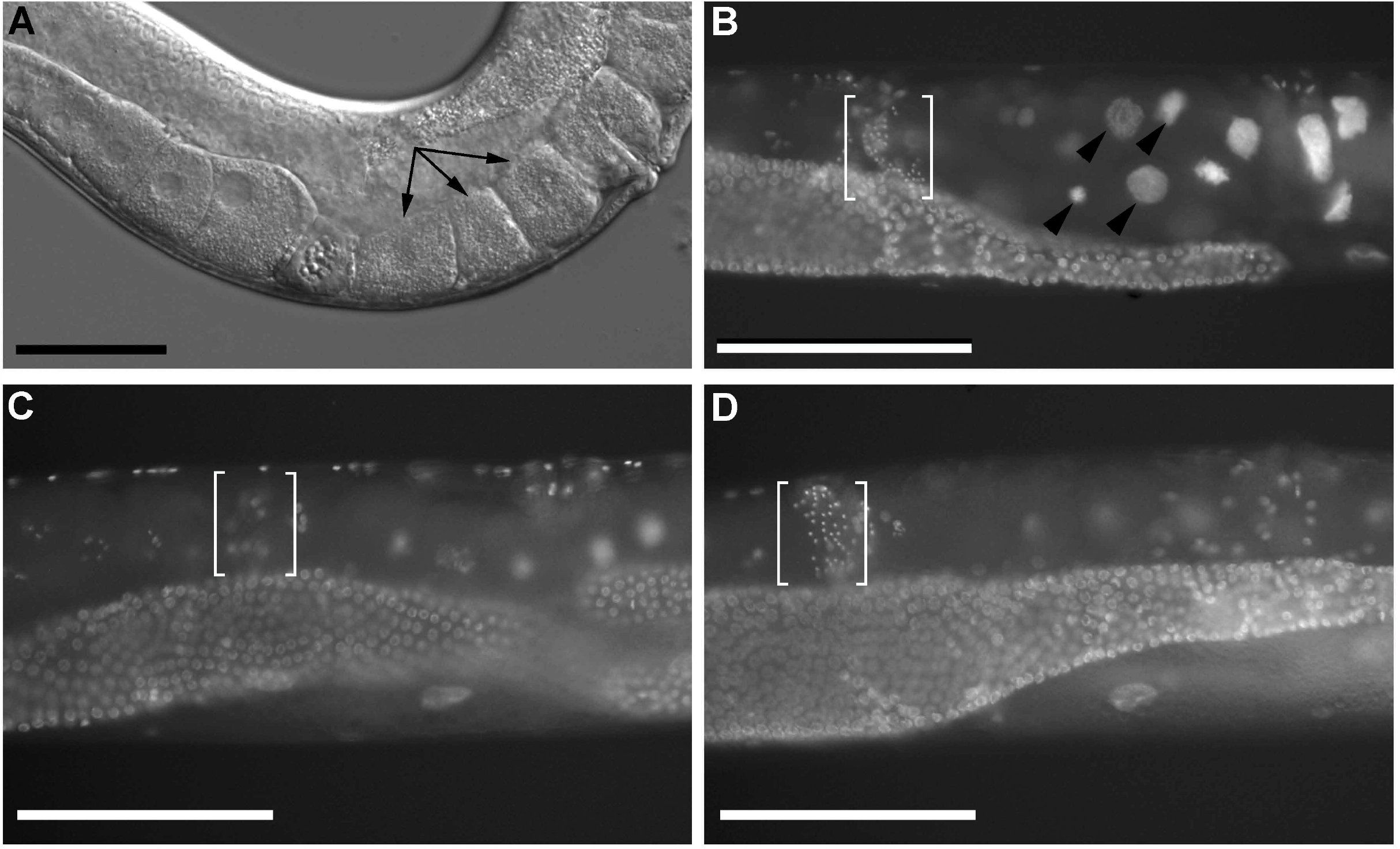
*spe-36* sperm are present but do not fertilize the eggs. (A) DIC imaging of *spe-36(as6)* hermaphrodites shows that oocytes are present in the oviduct and sperm are present in the spermatheca. However unfertilized eggs (arrows) are present in the uterus instead of developing embryos, despite the fact that the sperm must have contacted the eggs when the eggs moved through the spermatheca (brackets) to enter the uterus. (B) DAPI staining confirms the presence of sperm in the spermatheca (small bright spots within brackets) of 1 day adult hermaphrodites. Single DNA masses in the eggs in the uterus (arrowheads) are consistent with a lack of sperm entry into the egg. (C-D) DAPI staining of 4 day adult hermaphrodites. By day 4 of adulthood wild-type hermaphrodites (C) have depleted their self-sperm through fertilization of their eggs. In contrast, *spe-36* hermaphrodites of the same age (D) still have a full complement of sperm in their spermatheca. This indicates that the mutant sperm do not fertilize and are capable of maintaining their position in the spermatheca. Scale bars = 50 µm

### *spe-36* males mate, transfer sperm, and their sperm migrate to the spermathecae

Sperm are present in the spermatheca of *spe-36* hermaphrodites and these sperm make contact with the oocyte but fail to fertilize, as indicated by the rows of unfertilized oocytes within the uteri of unmated *spe-36* hermaphrodites (Figure 2A-B). To confirm that *spe-36* male sterility is also attributable to the sperm fertilization defect, we assayed the ability of males to transfer sperm to hermaphrodites and the ability of those sperm to migrate to the spermathecae. We expressed CYLC-2 tagged with mNeonGreen to mark the sperm of both *spe-36(as6); him-5* males and control males (Krauchunas et al., 2020) and mated them with N2 hermaphrodites. When the hermaphrodites were examined the following day, they all had male sperm in and near the spermathecae (Figure 3). From these data we infer that *spe-36* males mate and transfer sperm and those sperm are able to migrate to the correct location in the hermaphrodite reproductive tract. Therefore, we conclude that sperm made by either sex specifically require *spe-36* for fertilization.

**Figure 3.**
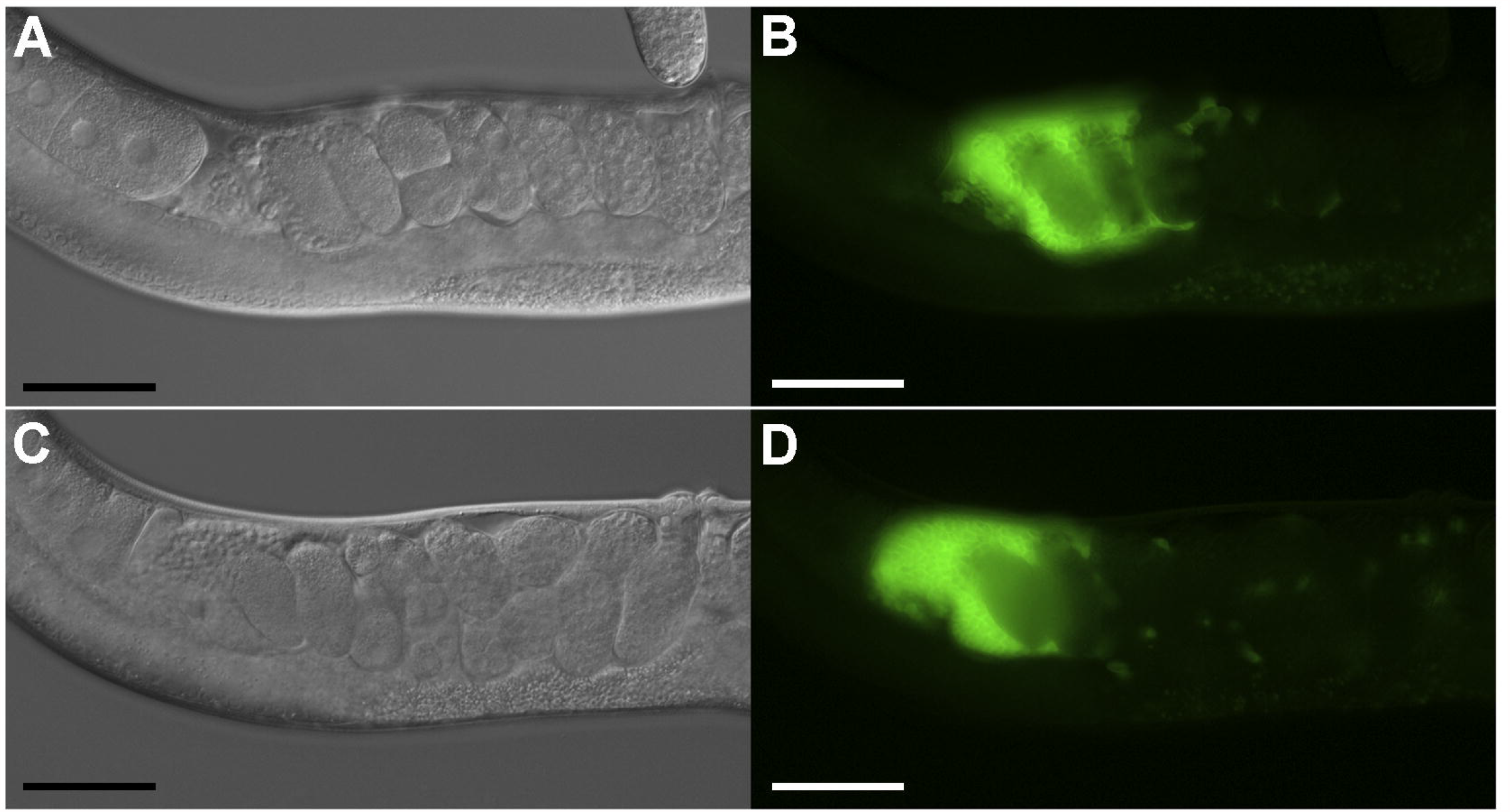
*spe-36* males exhibit normal mating, sperm transfer and sperm migratory behavior. *cylc-2 (mon-2[c41g7*.*6::mNG^3xFlag])* was crossed into *him-5* and *spe-36(as6); him-5*. Males with mNeonGreen (mNG) marked sperm were crossed with N2 L4 hermaphrodites and hermaphrodites were imaged ∼24 hours later. Both wild-type and mutant sperm were transferred to hermaphrodites and successfully migrated to the region of the spermatheca. (A- B) N2 hermaphrodite mated to *him-5; cylc-2::mNG* males (C-D) N2 hermaphrodite mated to *spe-36(as6); him-5; cylc-2::mNG* males. Scale bars = 50 µm

### Sperm from *spe-36* mutants are indistinguishable from wild-type sperm

To determine whether the fertility defects exhibited by *spe-36* worms are due to abnormal sperm morphology, we closely compared *spe-36* sperm to wild-type sperm. When examined using DIC optics, spermatids from *spe-36* mutant males were indistinguishable from wild-type spermatids (Figure 4A-B). Notably, spermiogenesis, the maturation of spherical, sessile spermatids into polar, motile spermatozoa (Muhlrad and Ward, 2002; Shakes and Ward, 1989), was unaffected; *spe-36* sperm activated normally *in vitro* (Figure 4D). When examined using transmission electron microscopy (TEM), spermatozoa within the reproductive tract of adult *spe-36* hermaphrodites also exhibited wild-type morphology (Figure 4E-F).

**Figure 4.**
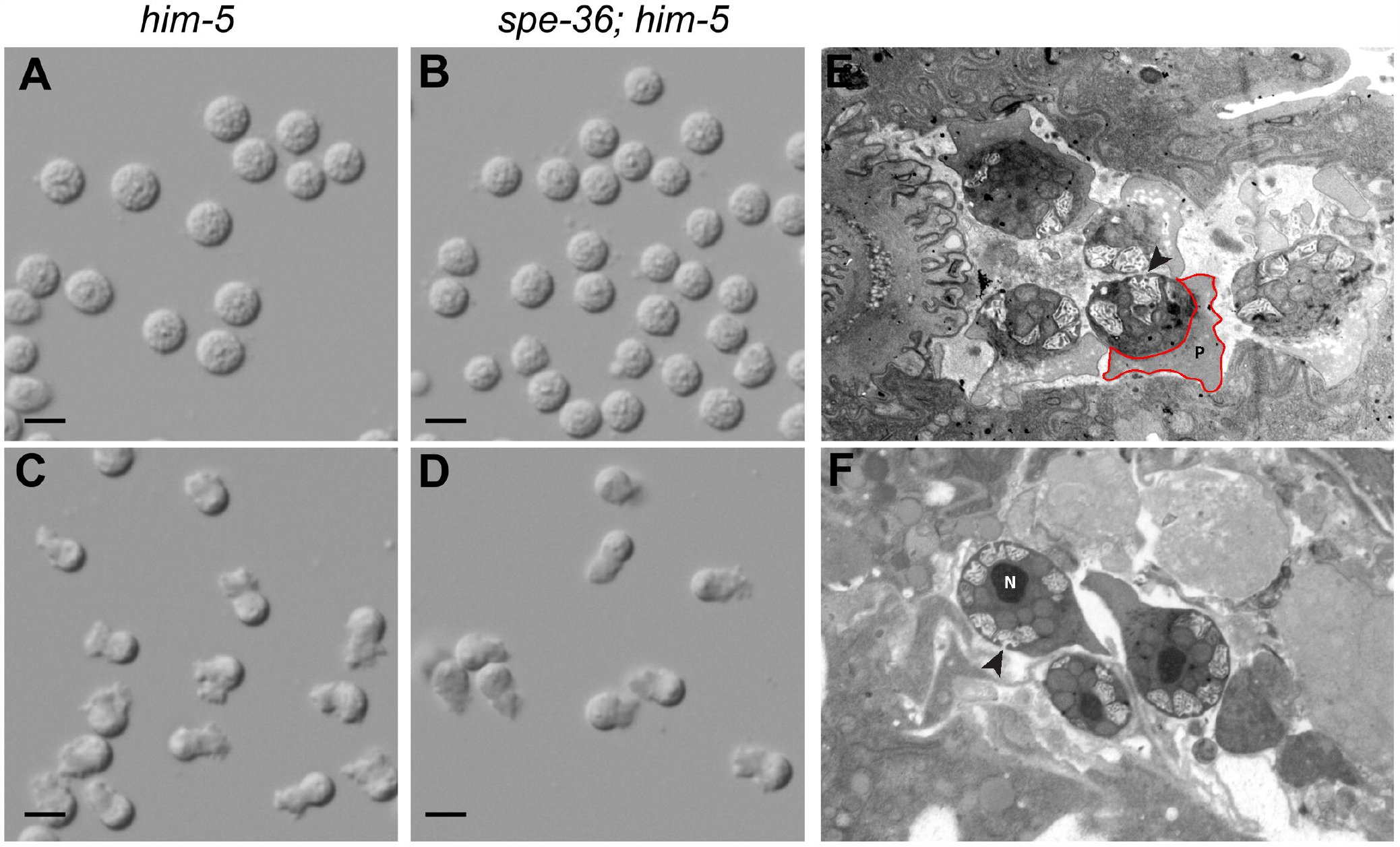
The morphology of spe-36 mutant sperm is indistinguishable from that of wild-type sperm. (A) *him-5* spermatids (B) *spe-36(as6); him-5* spermatids (C) Pronase activated *him-5* spermatozoa (D) Pronase activated *spe-36(as6); him-5* spermatozoa (E-F) Transmission electron micrographs of *spe-36(it114) unc-22* spermatozoa. The ultrastructural details of *spe-36* sperm are indistinguishable from those of wild type, including the sperm chromatin mass (N), the pseudopod (P), and the fused membranous organelles (arrowheads). Scale bars = 5 µm

### The *spe-36* gene encodes a protein with a single EGF-motif

We took a mapping-by-sequencing approach to determine the genomic locus of *spe-36* (Doitsidou et al., 2010). This strategy identified an approximately 3 Mbp region on chromosome IV (10,000,000 – 13,000,000 bp) that showed a strong reduction in Hawaiian SNP frequency. This region matched the predicted location of *spe-36(it114)* based on traditional mapping data (data not shown) making it the target region for further analysis of the whole genome sequencing data. We sequenced both *spe-36(it114)* and *spe-36(as6)* strains which allowed us to filter for common genes between the two data sets that had unique homozygous single nucleotide polymorphisms (SNPs). This analysis produced *F40F11*.*4* as the only likely candidate for *spe-36*. We identified a missense mutation (C104Y) in *F40F11*.*4* in *spe-36(it114)* and a splice donor variant (G ->A) in the first intron in *F40F11*.*4* in *spe-36(as6)* (Figure 1B). Sanger sequencing confirmed that neither of these SNPs are present in our lab N2 strain and revealed that *spe-36(as1)* contains the same missense mutation as *spe-36(it114)*.

We created transgenic animals carrying extrachromosomal arrays containing a PCR product of the full *F40F11*.*4* genomic sequence (plus upstream and downstream sequence to the boundaries of neighboring genes) and tested for the ability to rescue *spe-36* sterility. We measured *spe-36(as6)* self-broods for two independent lines carrying the arrays and found that both were able to restore hermaphrodite fertility (Figure 1 and data not shown). From the sequencing and rescue data, we assign the name *spe-36* to *F40F11*.*4*.

The *spe-36* gene produces a single transcript that encodes a protein of 350 amino acids. Analysis of the protein sequence predicts a single epidermal growth factor (EGF)-like domain near the C-terminus. We were surprised to find that *spe-36* is not predicted to encode a transmembrane domain, but does contain a signal sequence. There is also no evidence that SPE-36 is a GPI-anchored protein (PredGPI). Until now, all identified sperm function proteins have at least one transmembrane domain (Krauchunas et al., 2016; Mei and Singson, 2021). This is expected for proteins whose role is to mediate adhesion and fusion between the plasma membrane of the sperm and the plasma membrane of the egg. However, the SPE-36 protein sequence suggests that SPE-36 is secreted.

### SPE-36 is secreted

To test our prediction that SPE-36 is a secreted protein we expressed SPE-36 tagged with GFP in body wall muscle cells and looked for uptake in the coelomocytes via endocytosis. *C. elegans* coelomocytes are scavenger cells that actively endocytose fluid and macromolecules from the body cavity (Fares and Grant, 2002). Thus, proteins secreted from the muscle will end up in the coelomocytes while non-secreted proteins present in the muscle will not. When expressed under the *myo-3* promoter, SPE-36::GFP is visible in the coelomocytes in addition to the body wall muscle (Figure 5C-D and S2), consistent with SPE-36 being a secreted protein. In contrast, the single-pass transmembrane protein SPE-9 tagged with GFP and expressed under the *myo-3* promoter is seen only in muscle cells (Figure 5E-F and S2).

**Figure 5.**
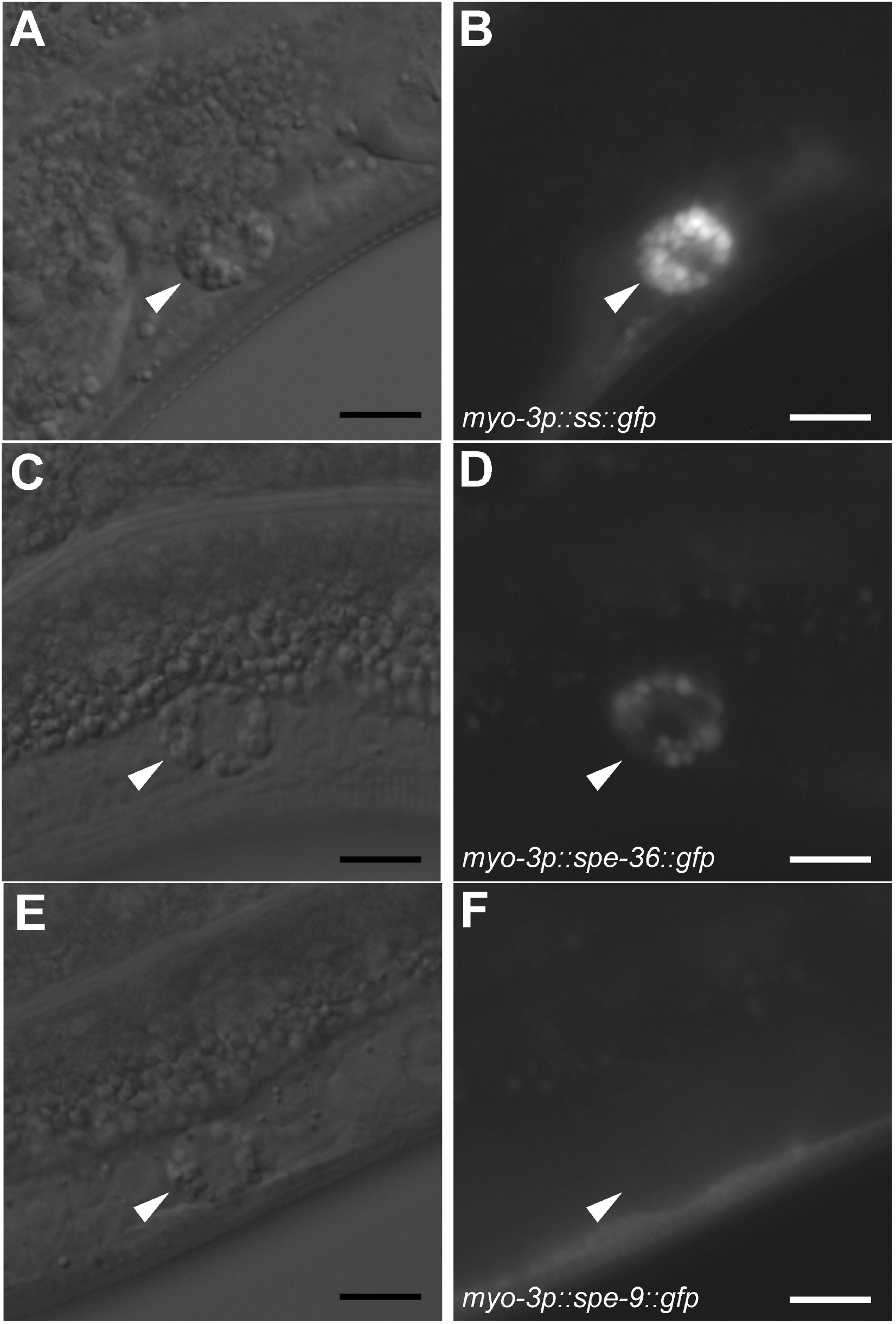
SPE-36 is a secreted protein. GFP-tagged proteins that are secreted from muscle cells can be taken up by the coelomocytes. When expressed under the *myo-3* promoter, SPE-36::GFP is present in coelomocytes (arrowheads) while the transmembrane protein SPE-9::GFP is not. (A-B) myo-3p::ss::gfp (C-D) myo-3p::spe-36::gfp (E-F) myo-3p::spe-9::gfp. Scale bars = 10 µm

### SPE-36 acts cell autonomously

Since SPE-36 is secreted we aimed to determine if it could act non-cell autonomously. To test this, we first crossed *spe-36(as6); him-5* males with *dpy-5* hermaphrodites. If secreted SPE-36 acts non-cell autonomously, we predicted that SPE-36 secreted by the hermaphrodite self sperm could render the *spe-36* male sperm functional and result in outcross progeny. However, the presence of wild-type hermaphrodite sperm did not increase the number of progeny sired by *spe-36* males (Table 1). Next, we tested whether SPE-36 secreted by male sperm could be used by *spe-36(as6)* hermaphrodite sperm to allow them to fertilize eggs. We crossed *spe-9(eb19); him-5* males, which are incapable of producing cross progeny due to the loss of SPE-9, with *spe-36(as6)* hermaphrodites to see if the hermaphrodites could now produce self-progeny. Again, we saw no increase in the number of progeny produced (Table 1). Thus, we found no evidence that *spe-36* mutant sperm can fertilize eggs when in the presence of SPE-36 producing sperm. While questions remain about the timing of SPE-36 secretion and action, our data strongly suggest that SPE-36 acts cell autonomously.

**Table 1.**
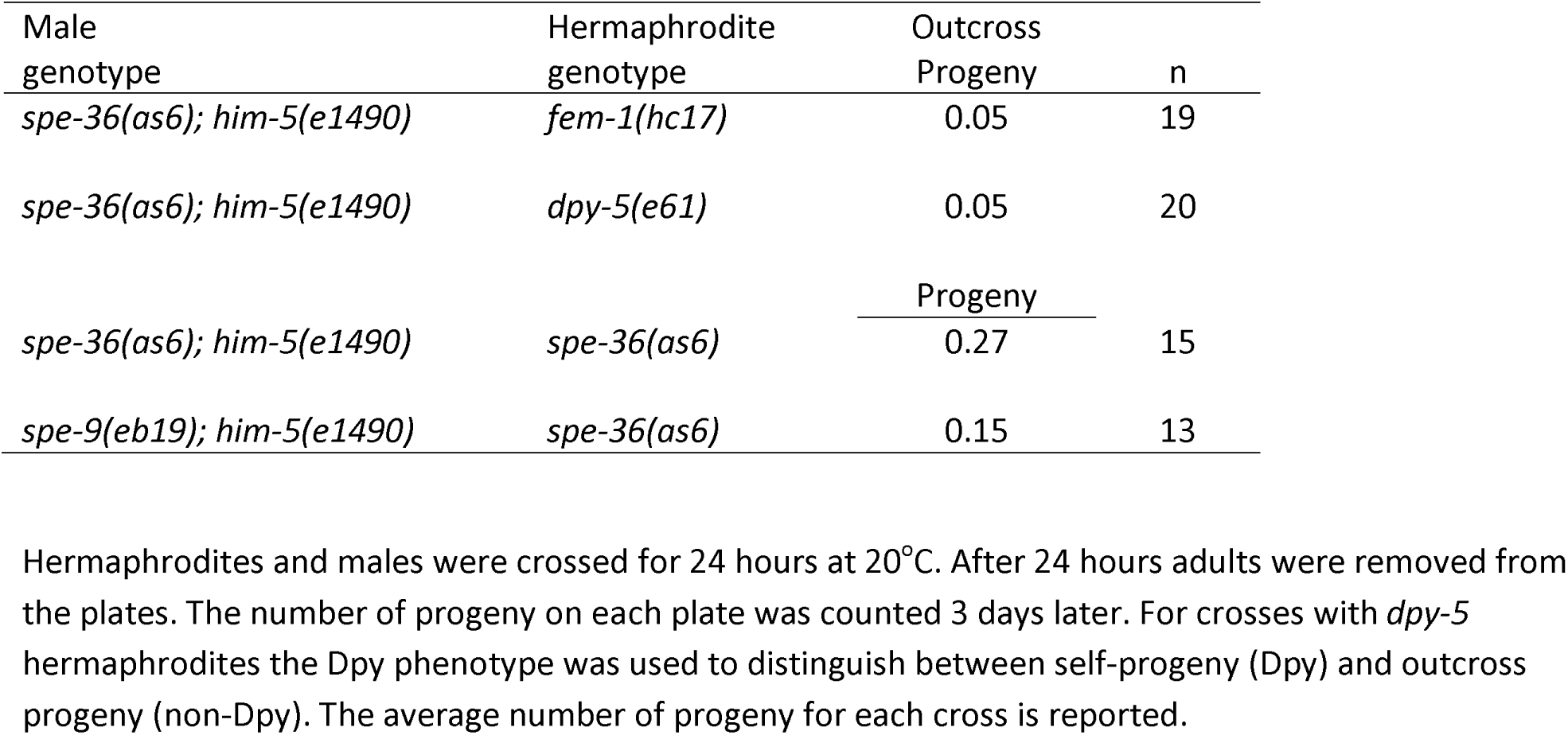
SPE-36 acts cell autonomously.

### SPE-36 is present in the cell body and on the pseudopod of activated spermatozoa

Our observation that SPE-36 acts cell autonomously suggests that SPE-36 remains localized to the sperm after it is secreted. To test this hypothesis, we used CRISPR to add a GFP tag to the C-terminus of SPE-36 at the endogenous locus. We were unable to see any GFP signal in live cells using either widefield or confocal microscopy (data not shown); therefore, we fixed isolated sperm and immunostained with an anti-GFP antibody. As SPE-36 functions during fertilization, we first observed its localization in activated spermatozoa. Sperm from *C. elegans* males go through spermiogenesis, i.e. sperm activation, when they are transferred to the hermaphrodite during copulation (Ward and Carrel, 1979). In wild-type *C. elegans*, sperm become fertilization-competent when they are activated (Ellis and Stanfield, 2014; Minniti et al., 1996). We were concerned that *in vitro* activators such as Pronase could have a negative effect on surface proteins and antibody binding. To look at SPE-36::GFP in *in vivo* activated sperm *spe-36::gfp; him-5* males were allowed to mate with *spe-36::gfp; him-5* hermaphrodites for 24 hours and then the hermaphrodites were dissected to release a combination of male and hermaphrodite spermatozoa. Anti-GFP antibody staining revealed a clear signal present in both the cell body and the along the leading edge of the pseudopod that is absent in sperm from *him-5* controls (Figure 6). The signal in the cell body often appears punctate and may represent SPE-36 that remains trapped in the membranous organelles (MOs) after MO fusion. All together our data are consistent with a model in which SPE-36 is localized to the MOs during spermatogenesis and released when the MOs fuse with the plasma membrane during sperm activation. After it is released, SPE-36 remains associated with the sperm surface on both the cell body and pseudopod where it can participate in binding or fusion with the egg.

**Figure 6.**
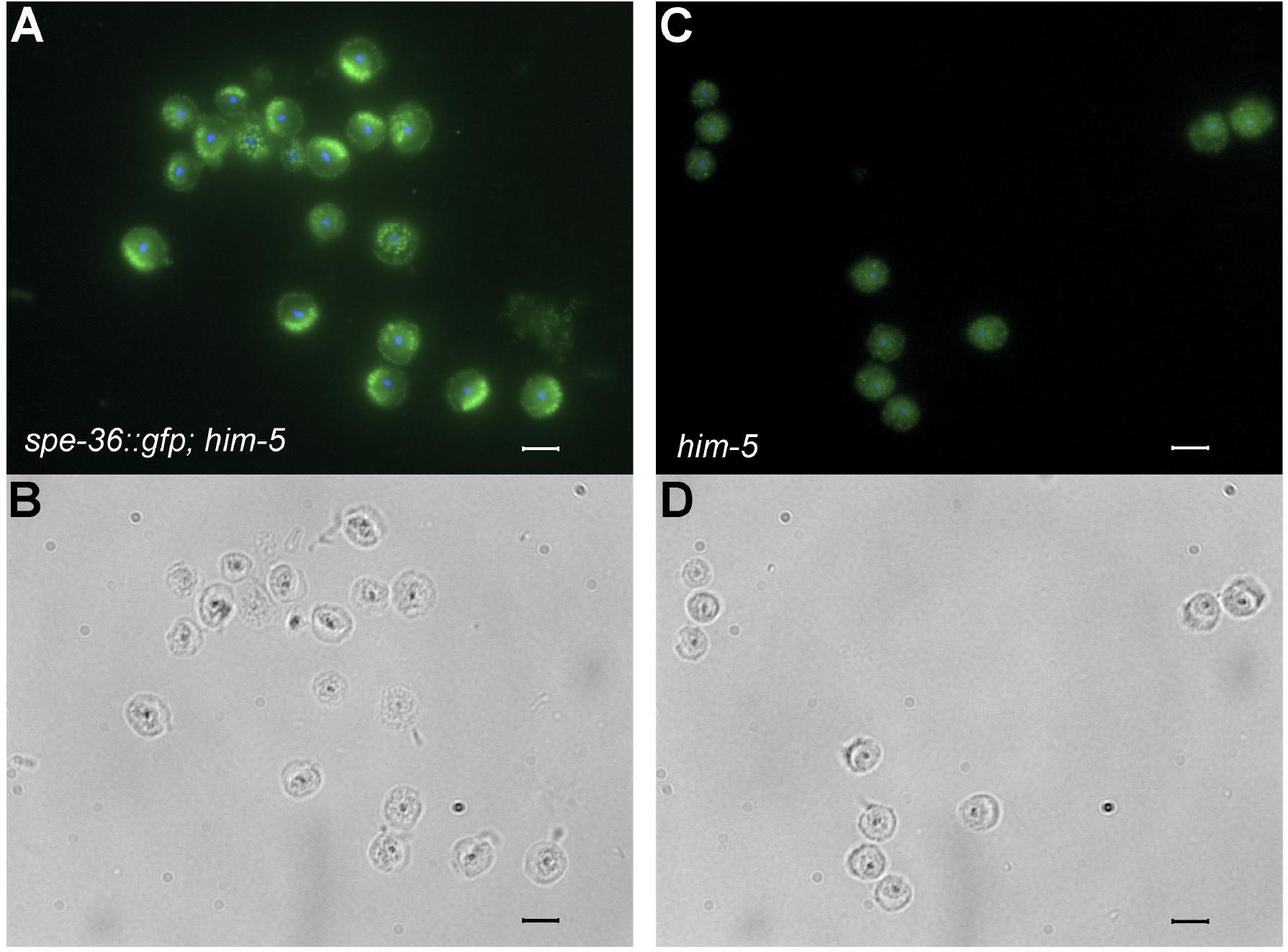
SPE-36 is localized to the cell body and pseudopod of activated sperm. *In vivo* activated spermatozoa from either *spe-36::gfp; him-5* (A-B) or *him-5* (C-D) animals were fixed and stained with an anti-GFP antibody. A signal is seen in both the cell body and along the edge of the pseudopod in sperm with SPE-36::GFP that is absent in the *him-5* controls. Scale bars = 5 µm

## Discussion

The fertilization synapse model proposes that specialized zones of interaction and multi- protein complexes composed of both *trans* and *cis* protein-protein interactions mediate gamete interaction and fusion. This model accounts for the likely presence of molecules that mechanistically ensure adhesion and fusion, as well as additional molecules that provide species-specific recognition between the gametes. The number of proteins necessary for fertilization that contain domains utilized for protein-protein interactions further argue for multi-protein complex(es) present on the gametes that mediate gamete interaction and fusion.

Our discovery of SPE-36 adds a new class of proteins to our model of the fertilization synapse. All previously discovered sperm function genes encode single- and multi-pass transmembrane proteins, which matches our expectations about the types of proteins that regulate membrane interactions. SPE-36 is a secreted protein that is produced by sperm and functions cell-autonomously to mediate fertilization. The *spe-36* mutants phenocopy other known ‘sperm-function’ fertility mutants (reviewed in Mei and Singson, 2021). Mutant *spe-36* sperm develop normally and ultimately mature into sperm that are morphologically indistinguishable from wild type at the level of both light and electron microscopy. Sperm from *spe-36* mutants are fully motile and can both migrate to and maintain their position within the spermatheca. Yet, *spe-36* mutant sperm are incapable of fertilizing wild-type oocytes regardless of whether the sperm are hermaphrodite or male derived and despite having frequent contact with mature oocytes in the spermatheca.

BLAST searches show no obvious SPE-36 homologs outside of nematodes and some insects. Even though there may not be direct conservation of protein sequence, we predict organisms may use a similar set of protein domains to mediate fertilization. For example, BLAST searches do not identify *C. elegans* SPE-45 and mammalian IZUMO1 as homologs. However, both proteins contain similar domain structures and loss-of-function mutations result in the same sterile phenotype in worms and mice, respectively (Nishimura et al., 2015; Singaravelu et al., 2015). Likewise, the gamete fusogen HAP2/GCS1 is incredibly similar in protein structure to viral class II fusogens despite a lack of similarity between their primary amino acid sequences (Clark, 2018; Fedry et al., 2018; Fédry et al., 2017; Pinello et al., 2017; Valansi et al., 2017).

We are beginning to find that there are multiple proteins with shared domains among the collection of sperm function proteins. In *C. elegans*, both SPE-45 and SPE-51, contain an Ig-like domain (Nishimura et al., 2015; Singaravelu et al., 2015; Mei et al., co-submission). Similarly, both IZUMO1 and SPACA6 are proteins with Ig-like domains that are required for mammalian fertilization (Barbaux et al., 2020; Inoue et al., 2005; Lorenzetti et al., 2014; Noda et al., 2020). Fertilization in both worms and mammals also requires a pair of DC-STAMP domain-containing proteins: SPE-42 and SPE-49 in *C. elegans*, DCST1 and DCST2 in mammals (Inoue et al., 2021; Kroft et al., 2005; Noda et al., 2021; Wilson et al., 2018). Mammalian ADAM3, Integrin Beta, SED1, and *C. elegans* SPE-9 and SPE-36, all contain one or more EGF domains (Ensslin and Shur, 2003; Singson et al., 1998; Takagi et al., 2001; Yuan et al., 1997). EGF motif containing proteins are commonly found on the cell surface and are known to mediate cell adhesion in other contexts (Carpenter and Cohen, 1990; Stenflo, 1991; Wagner and Wyss, 1994). Thus, the presence of EGF motifs in these proteins suggests a potential role in cell adhesion and important protein-protein interactions during fertilization. There is also evidence that interactions can take place between these different types of domains that exist in sperm function genes. For example, during neuronal synapse development the interaction of an EGF domain containing protein, Caspr4, and an Ig domain containing protein, NB2, on the cell surface is necessary to stabilize the synapse (Ashrafi et al., 2014).

The localization of SPE-36 and cell autonomous nature of SPE-36 function show that the protein remains associated with the sperm membrane after it is secreted. Thus, we predict that *cis*-interactions with other sperm function proteins keep SPE-36 tethered to the sperm. Whether SPE-36 is important for the assembly of a multi-protein complex on the sperm surface or directly interacts with one or more proteins on the egg surface remains to be determined. Genetic and biochemical experiments will be needed to identify the specific proteins that interact with SPE-36. Our studies suggest that gamete expressed genes that encode proteins with EGF motifs as well as secreted proteins should be investigated for roles in fertilization in other species. As we continue with candidate gene approaches (Holcomb et al., 2020; Lu et al., 2019; Miyata et al., 2016; Noda et al., 2019) we need to realign our expectations of the types of proteins that represent viable candidates. As we work out the *cis*-interactions at the surface of each gamete, and the *trans*-interactions between the sperm and egg, we continue to grow our model of a fertilization synapse that goes far beyond the recognition between a ligand and receptor pair.

## Acknowledgments

We would like to thank Diane Shakes for the original discovery of *spe-36(it114)*, Steve L’Hernault for confirming the phenotype, Pavan Kadandale for assistance mapping the allele, David Hall for assistance with electron microscopy, and Jeremy Bird for careful reading of the manuscript. This work was supported by National Institutes of Health R01HD054681 to AS. We thank the CGC (which is funded by NIH Office of Research Infrastructure Programs P40 OD010440) and WormBase.

## Author Contributions

Conceptualization, A.R.K., M.R.M., X.M., and A.S.; Methodology, A.R.K., M.R.M., X.M., and A.S.; Investigation, A.R.K., M.R.M., A.L., E.P., G.S., and I.I.A.; Writing – Original Draft, A.R.K. and M.R.M.; Writing – Review & Editing, A.R.K., M.R.M., X.M., and A.S.; Supervision, A.R.K., M.R.M., and A.S.; Funding Acquisition, A.S.

---

**Figure S1.**
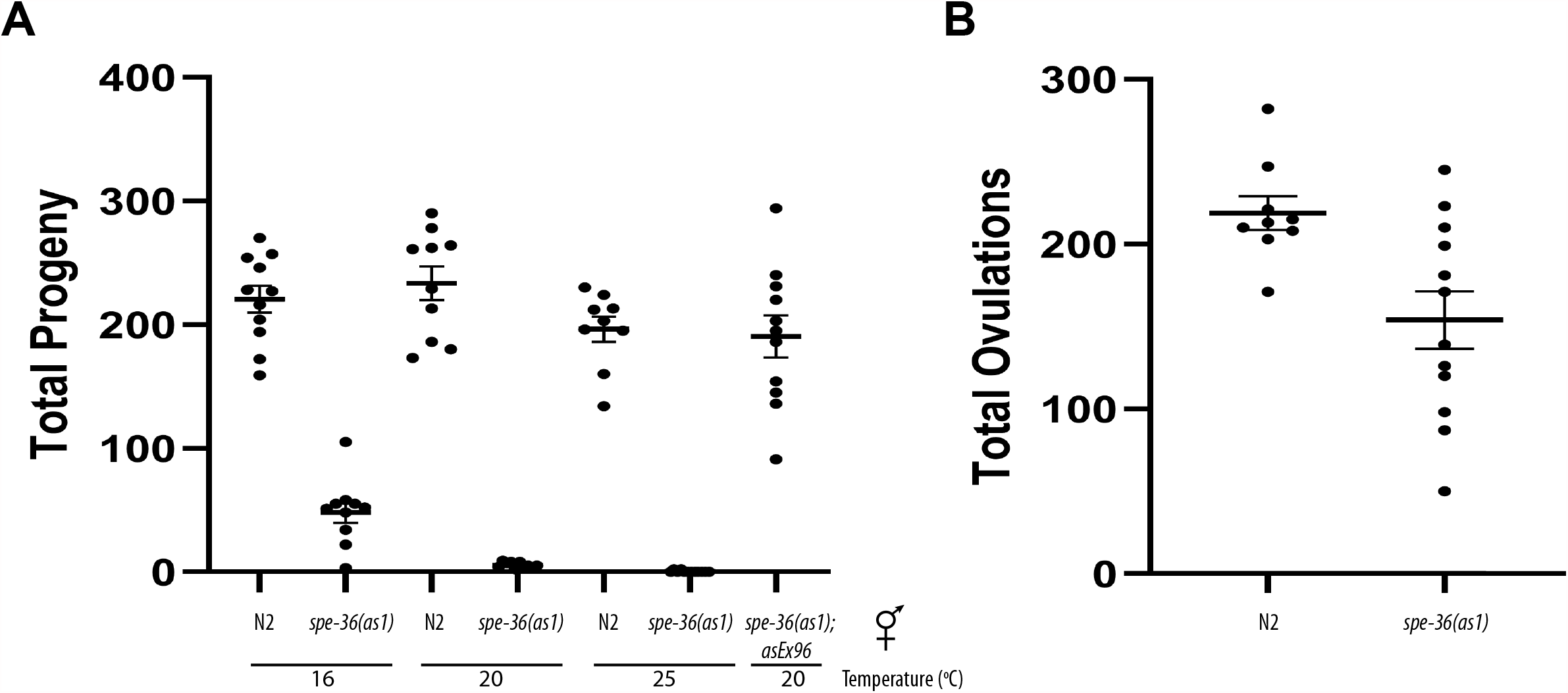
*spe-36(as1)* mutants display a temperature-sensitive phenotype but continue to ovulate at high rates at the non-permissive temperature. Even at lower temperatures *spe- 36(as1)* hermaphrodites produce a highly reduced number of progeny relative to wild-type controls.

**Figure S2.**
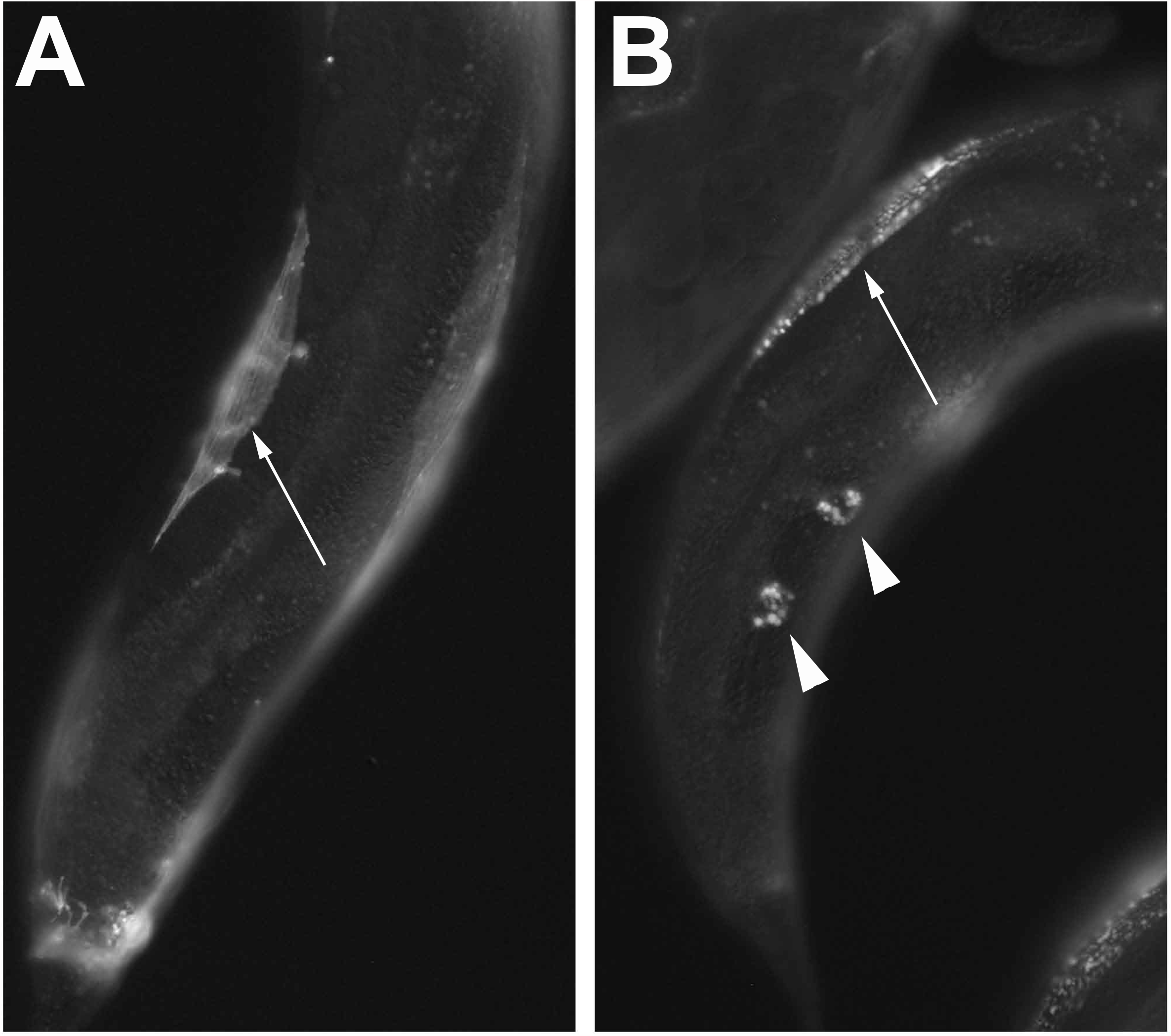
Both SPE-9::GFP (A) and SPE-36::GFP (B) can be seen in the body wall muscle cells (arrows) when expressed under the *myo-3* promoter. However only SPE-36::GFP is seen in the coelomocytes (arrowheads) indicating that SPE-36 is secreted while SPE-9 is not.

